# Genetic characterization of cucumber genetic resources in the NARO Genebank indicates their multiple dispersal trajectories to the East

**DOI:** 10.1101/2024.04.28.591496

**Authors:** Gentaro Shigita, Koichiro Shimomura, Dung Phuong Tran, Naznin Pervin Haque, Thuy Thanh Duong, Odirich Nnennaya Imoh, Yuki Monden, Hidetaka Nishida, Katsunori Tanaka, Mitsuhiro Sugiyama, Yoichi Kawazu, Norihiko Tomooka, Kenji Kato

## Abstract

The cucumber (*Cucumis sativus*) is an economically important vegetable crop cultivated and consumed worldwide. Despite its popularity, the manner in which cucumbers were dispersed from their origin in South Asia to the rest of the world, particularly to the east, remains a mystery due to the lack of written records. In this study, we performed genotyping-by-sequencing (GBS) on 723 worldwide cucumber accessions, mainly deposited in the Japanese National Agriculture and Food Research Organization (NARO) Genebank, to characterize their genetic diversity, relationships, and population structure. Analyses based on over 60,000 genome-wide single-nucleotide polymorphisms identified by GBS revealed clear genetic differentiation between Southeast and East Asian populations, suggesting that they reached their respective region independently, not progressively. A deeper investigation of the East Asian population identified two subpopulations with different fruit characteristics, supporting the traditional classification of East Asian cucumbers into two types thought to have been introduced by independent routes. Finally, we developed a core collection of 100 accessions representing at least 93.2% of the genetic diversity present in the entire collection. The genetic relationships and population structure, their associations with geographic distribution and phenotypic traits, and the core collection presented in this study are valuable resources for elucidating the dispersal history and promoting the efficient use and management of genetic resources for research and breeding in cucumber.

**Key message:** Genotyping-by-sequencing of 723 worldwide cucumber genetic resources revealed that cucumbers were dispersed eastward via at least three distinct routes, one to Southeast Asia and two from different directions to East Asia.

## Introduction

The cucumber (*Cucumis sativus*) is one of the most economically important vegetable crops grown and consumed worldwide, with an agricultural production covering two million hectares and a yield of over 90 million tons of fruit annually (FAOSTAT 2022; https://www.fao.org/faostat). The cultivated cucumber (*C*. *s*. var. *sativus*) was domesticated about 3,000 years ago in the Himalayan foothills from its wild form, *C*. *s*. var. *hardwickii*, which produces small, bitter, seedy fruits (Bisht et al. 2004; Chomicki et al. 2020; Renner et al. 2007; Sebastian et al. 2010). From its origins in the Indian subcontinent, the cucumber appears to have traveled both westward and eastward, following several trajectories in each case (Meglic et al. 1996; Staub et al. 2002; Staub et al. 1999). To the west, iconographic and lexicographic evidence suggests that cucumbers had arrived in Europe by the second half of the tenth century via two independent routes, overland and maritime, and subsequently spread to Africa and the Americas (Paris et al. 2012). To the east, although there is written evidence that cucumbers had been introduced to northern China from the west through the Silk Road by about 2,000 years ago (Li 1969; Li 1590), less is known about when and by what route(s) they reached southern China and Southeast Asia.

Attempts to investigate the genetic diversity of the cucumber began with a series of studies using the genetic resource collection deposited in the U.S. National Plant Germplasm System (NPGS) based on isozymes (Knerr and Staub 1992; Knerr et al. 1989; Meglic et al. 1996; Meglic and Staub 1996; Staub et al. 1999; Staub et al. 1997) or random amplified polymorphic DNA (RAPD) markers (Horejsi and Staub 1999). These studies led to the selection of a small subset of accessions, the so-called core collection, representing morphological, geographic, and genetic diversity (Staub et al. 2002). The study with the most extensive sampling to date employed 3,342 accessions compiled from several national genetic resource centers (Lv et al. 2012). Analysis of this mega-collection with 23 highly polymorphic simple sequence repeat (SSR) markers revealed three distinct genetic populations that largely corresponded to three geographic regions: 1) East Asia excluding the Xishuangbanna Dai Autonomous Prefecture (hereafter Xishuangbanna) in tropical southwestern China; 2) regions west of India, such as Europe, the Americas, and West and Central Asia; and 3) India and Xishuangbanna. This study eventually led to the construction of a core collection consisting of 115 accessions that cover 77.2% of the SSR alleles found in the mega-collection. Deep resequencing of these 115 accessions identified over three million variations and further distinguished a population endemic to Xishuangbanna from Indian accessions (Qi et al. 2013). This population has served as another botanical variety, *C*. *s*. var. *xishuangbannanensis*, due to its unique characteristics, such as thick cylindric fruits with five or more carpels and orange flesh resulting from β-carotene accumulation (Qi et al. 1983; Renner 2017). The genetic–geographic grouping of cucumbers has been generally confirmed by subsequent studies with varying sets of genetic resources (Lee et al. 2020; Liu et al. 2019a; Wang et al. 2018). Ultimately, a graph-based pan-genome was constructed from chromosome-scale genome assemblies of 12 representative accessions, facilitating the comprehensive identification of genetic variations, including large chromosomal rearrangements and structural variations associated with agronomic traits (Li et al. 2022).

However, despite the numerous genetic resources used in these studies, little attention has been paid to cucumbers in Southeast Asia. For example, only 29 accessions from Southeast Asian countries were included in the 3,342 accessions studied by Lv et al. (2012). Similarly, only 11 of the 1,234 accessions examined by Wang et al. (2018) were of Southeast Asian origin. This is probably due to the limited availability of cucumber accessions from Southeast Asia in major national genetic resource centers, such as the U.S. NPGS, the National Crop Genebank of China (NCGC), the Centre for Genetic Resources, the Netherlands (CGN), and the Leibniz Institute of Plant Genetics and Crop Plant Research (IPK). As of March 2024, the U.S. NPGS holds over 2000 cucumber accessions, of which only 16 are from Southeast Asian countries. This underrepresentation of Southeast Asian cucumbers raises concerns that the previous studies listed above may have underestimated genetic diversity or even overlooked another distinct genetic population present in Southeast Asia. In this context, the cucumber collection deposited in the Japanese genetic resource center, the National Agriculture and Food Research Organization (NARO) Genebank, could provide the opportunity to fill this potential gap and track the eastward dispersal trajectories of cucumber, as it contains a number of genetic resources recently introduced from four Southeast Asian countries: Cambodia (Kawazu et al. 2020; Matsunaga et al. 2015; Tanaka et al. 2016; Tanaka et al. 2019; Tanaka et al. 2017; Yashiro et al. 2019); Laos (Okuizumi et al. 2015); Myanmar (Shimomura et al. 2020); and Vietnam (Kawazu et al. 2017; Shimomura et al. 2016; Sugiyama et al. 2015), through collaborative expeditions conducted under the Plant Genetic Resources in Asia (PGRAsia) project (https://sumire.gene.affrc.go.jp/pgrasia/index_en.php).

In this study, we conducted genetic characterization of 723 worldwide cucumber accessions, mainly deposited in the NARO Genebank, using the genotyping-by-sequencing (GBS) approach (Elshire et al. 2011). Phylogenetic and population structure analyses based on the identified genome-wide single-nucleotide polymorphisms (SNPs) revealed four distinct genetic populations highly consistent with their geographic distributions. These populations exhibited differing characteristics in various phenotypic traits, which may reflect the effects of artificial selection during their dispersal to each geographic region. Furthermore, our in-depth analysis of the East Asian population identified two subpopulations that differ in specific fruit traits, supporting the traditional classification of East Asian cucumbers into two types believed to have been introduced via independent routes. Finally, we developed a core collection of 100 accessions covering most of the genetic diversity present in the entire collection to facilitate the efficient use and management of genetic resources for research and breeding in cucumber.

## Materials and Methods

### Plant materials

A total of 723 cucumber accessions were used in this study, comprising 708 accessions deposited in the NARO Genebank and 15 commercial cultivars. These accessions originated from diverse geographic regions: 352 from East Asia, 118 from Europe, 107 from Southeast Asia, 93 from South Asia, 34 from the Americas, 14 from Central and West Asia, three from Oceania, and two from Africa. The materials included one accession of *C*. *s*. var. *hardwickii* from Nepal. In addition, two accessions of *C. hystrix*, a sister species of cucumber (Endl et al. 2018; Renner et al. 2007; Sebastian et al. 2010), were used as the outgroup. Detailed information on all accessions used is given in Table S1.

### Genotyping-by-sequencing

GBS library preparation and sequencing followed the same procedure as Shigita et al. (2023). In brief, the genomic DNA of each accession was extracted from young leaves using the cetyltrimethylammonium bromide-based method (Murray and Thompson 1980) with minor modifications. The DNA was then digested with either *Hsp*92II (Promega, Madison, WI, USA) or *Nla*III (New England Biolabs, Ipswich, MA, USA) restriction enzyme. Adapters containing an 8-bp barcode sequence for sample identification were ligated to both ends of the restricted DNA fragments, and the resulting library was enriched by PCR amplification. Finally, the libraries of 192 accessions were pooled and sequenced on the HiSeq 4000 platform (Illumina, San Diego, CA, USA), generating 101-bp paired-end reads. The raw sequencing reads were sorted into individual accessions according to the combination of the 8-bp barcode sequences at both ends.

Adapter sequences and low quality bases were trimmed from the raw reads using fastp v0.20.0 (Chen et al. 2018) with the setting “-3 -l 20 -a=AGATCGGAAGAGC”. SNP calling was performed using the GB-eaSy pipeline (Wickland et al. 2017). First, the clean reads of each accession were mapped to the reference genome of the ‘Chinese Long’ inbred line 9930 v3.0 (Li et al. 2019), using BWA-MEM v0.7.17 (Li 2013) with default parameters. Next, the mpileup and call commands in BCFtools v1.17 (Danecek et al. 2021) were used to generate a pileup from the mapped reads and identify candidate SNPs. Finally, raw SNPs were called using VCFtools v0.1.16 (Danecek et al. 2011) with the parameters “--mac 1 --minDP 3”. Filtered SNPs were obtained by removing sites with excessive missing data from the raw SNPs.

### Phylogenetic and population structure analyses

The filtered SNPs with a missing rate of less than 90% were used for phylogenetic and population structure analyses. A maximum likelihood phylogenetic tree was inferred using IQ-TREE v2.2.0.3 (Minh et al. 2020) under the GTR model with ascertainment bias correction (Lewis 2001), with 1,000 ultrafast bootstrap replicates (Hoang et al. 2018). A population structure analysis was performed using ADMIXTURE v1.3.0 (Alexander et al. 2009). To determine an appropriate number of ancestral populations (*K*), the program was first run for *K* from 1 to 10, with 10-fold cross-validation (--cv=10). Ten independent runs were then performed for each *K* from 2 to 4, and the major mode of each *K* was visualized using pong v1.5 (Behr et al. 2016).

The unfiltered raw SNPs were used to calculate the nucleotide diversity (*π*) (Nei and Li 1979) and the population fixation index (*F*_ST_) (Weir and Cockerham 1984), using VCFtools v0.1.16 (Danecek et al. 2011). The average *π* for each population was calculated as the sum of *π* at each SNP site divided by the number of bases covered by GBS reads in the reference genome. The pairwise weighted *F*_ST_ values for each pair of populations were calculated and converted to two-dimensional coordinates by multidimensional scaling, using the cmdscale function in R v4.1.0.

### Phenotypic data compilation

To investigate the association of phenotypic traits with the genetic populations identified in the above analyses, evaluation data for 12 traits collected from cultivation trials conducted at 10 locations in Japan and Vietnam between 1983 and 2022 were downloaded from NARO’s Plant Genetic Resources Database (https://www.gene.affrc.go.jp/databases-plant_search_char_en.php) and summarized by accession. Of the 12 traits, three were quantitative (fruit length, fruit diameter, and seed length) and nine were qualitative (sex expression, number of female flowers per node, fruit bearing position, parthenocarpy, resistance to melon yellow spot virus, resistance to papaya ringspot virus, mature fruit skin color, mature fruit skin netting, and fruit spine color). If an accession had several inconsistent observations, we used the mean value for the quantitative traits, while it was considered as intra-accession segregation for the qualitative traits.

### Core collection selection

A core collection is a small subset of accessions that represents the genetic diversity present in a larger genetic resource collection. It plays a crucial role in facilitating the efficient use and management of genetic resources by providing researchers and breeders with a common, more tractable diversity panel to work with (Frankel and Brown 1984). To objectively select a core subset that represents the genetic diversity, accessions were ranked to maximize the cumulative allelic coverage of the filtered SNPs, using GenoCore (Jeong et al. 2017) with the setting “-cv 100”. The outgroup and 102 F_1_ accessions were excluded from this analysis. Additional accessions were manually selected based on their historical significance, geographic origin, or unique traits.

## Results

### Genome-wide SNP identification

The GBS yielded over 1.5 billion reads, for a total of 155 Gb of sequence data. The number of reads for each accession ranged from 137,950 to 16,768,608, with an average of 2,120,118 reads (Table S1). On average, 93.3% of the reads were mapped to the ‘Chinese Long’ inbred line 9930 genome (Li et al. 2019), covering 3.8% of the genome (8.7 Mb of 226.2 Mb) at a depth of 15.0 x. The GB-eaSy pipeline (Wickland et al. 2017) identified a total of 324,664 raw SNPs distributed throughout the genome with a high density of one SNP per 697 bp (Table 1). Retaining SNPs with a missing rate of less than 90% resulted in 61,367 filtered SNPs distributed at a density of one SNP per 3.7 kb.

**Table 1.** Summary statistics of the identified SNPs across cucumber chromosomes.

| Chromosome | Length (bp) | Raw SNPs | Average raw<br>SNP distance (bp) | Filtered SNPs<br>(missing rate < 90%) | Average filtered<br>SNP distance (bp) |
| --- | --- | --- | --- | --- | --- |
| Chr1 | 32,926,272 | 49,101 | 671 | 9,085 | 3,624 |
| Chr2 | 24,837,039 | 39,019 | 637 | 6,595 | 3,766 |
| Chr3 | 40,877,379 | 66,219 | 617 | 11,622 | 3,517 |
| Chr4 | 26,827,763 | 43,077 | 623 | 8,367 | 3,206 |
| Chr5 | 31,913,682 | 42,342 | 754 | 8,532 | 3,740 |
| Chr6 | 31,125,843 | 44,866 | 694 | 7,963 | 3,909 |
| Chr7 | 22,466,726 | 34,789 | 646 | 6,656 | 3,375 |
| Scaffolds | 15,236,958 | 5,251 | 2,902 | 2,547 | 5,982 |
| Overall | 226,211,662 | 324,664 | 697 | 61,367 | 3,686 |

### Phylogenetic relationships and population structure of the cucumber genetic resources

To infer the phylogenetic relationships among the 723 cucumber accessions, we inferred a maximum likelihood tree based on the filtered SNPs, using two accessions of *C. hystrix* as the outgroup. The phylogeny showed genetic relationships that largely corresponded to geographic distribution (Fig. 1a, Fig. S1). The *C*. *s*. var. *hardwickii* accession from Nepal (JP140500) was positioned closest to the outgroup and isolated from all other accessions by a long branch, confirming its wild status. Consistent with the South Asian origin of cucumber, the deepest branches near the root were occupied by accessions from South Asia. Accessions outside South Asia generally clustered into Southeast Asia, East Asia, and west of India, including Europe, the Americas, and Central and West Asia.

**Fig. 1.**
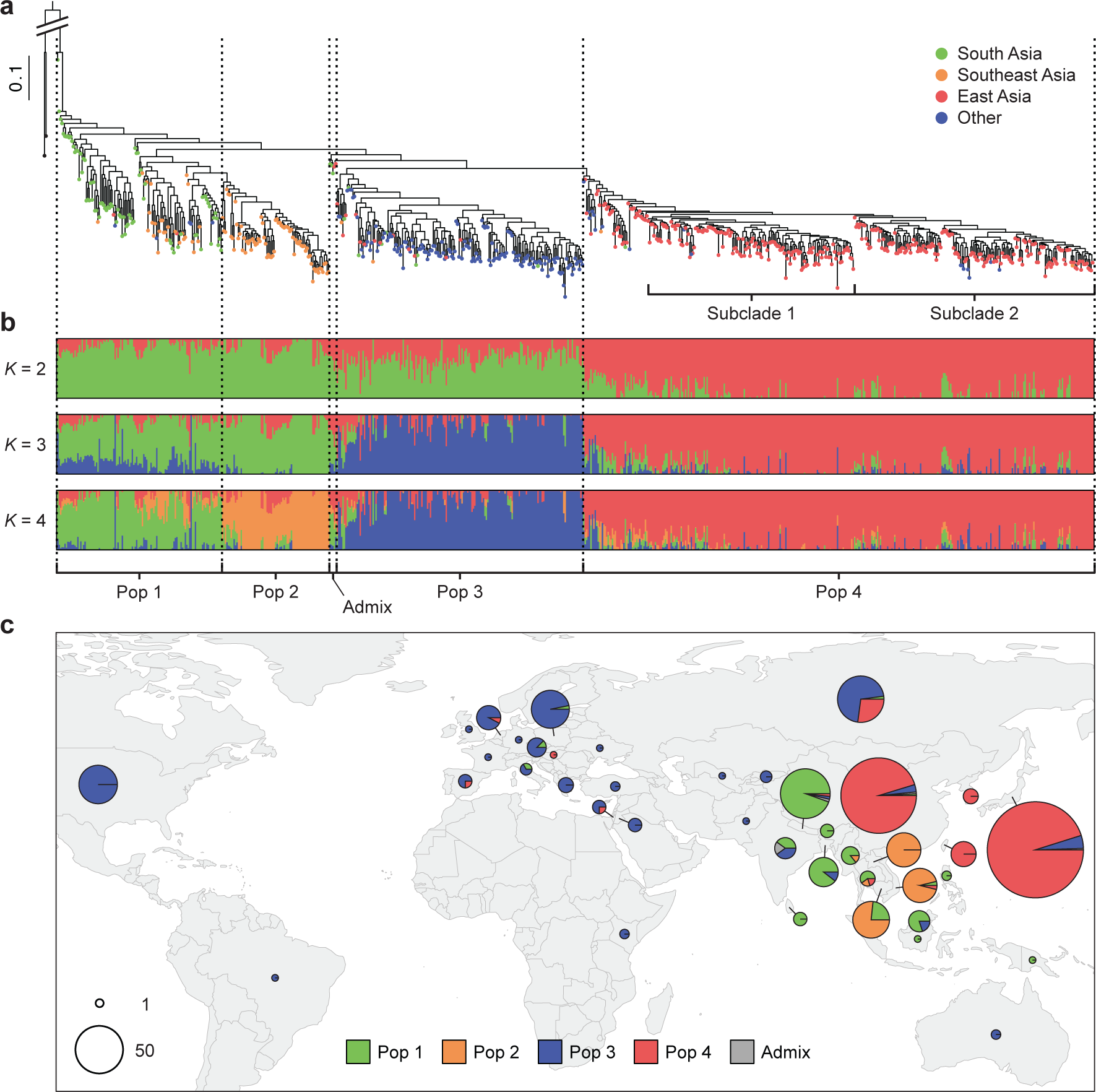
Phylogenetic relationships, population structure, and geographic distribution of the cucumber genetic resources. **a** Maximum likelihood phylogenetic tree of the 723 cucumber and two *C. hystrix* accessions inferred using the 61,367 filtered SNPs. Colors at the tips indicate the geographic origin of each accession. (A tree with individual tip labels and node supports is given in Fig. S1.) **b** Model-based clustering with *K* from 2 to 4. The *x*-axes list the accessions in the same order as they appear on the phylogenetic tree, and the *y*-axes quantify the membership to each ancestral population, represented in different colors. **c** Geographic distribution of the different genetic populations. For each country, the pie represents the proportion of the different populations, and its size reflects the relative sample size. The map was created using the rworldmap package (South 2011) in R v4.1.0

The genetic population structure of the 723 cucumber accessions was further investigated using the model-based clustering implemented in ADMIXTURE (Alexander et al. 2009). The preliminary runs for different values of *K* from 1 to 10 showed that the cross-validation error was minimized at *K* = 4 (Fig. S2), suggesting the presence of four primary genetic populations. More intensive runs were then performed for *K* values from 2 to 4 (Fig. 1b). At *K* = 2, accessions from East Asia were distinguished from all others. At *K* = 3, accessions west of India were identified as a separate population. At *K* = 4, a subset of Southeast Asian accessions was further distinguished as a distinct population. These results largely confirm the genetic differentiation of cucumbers linked to the geographic distribution observed in the phylogenetic analysis.

Taking the results of the phylogenetic and population structure analyses, we classified the 723 accessions into four genetic populations (Pops 1–4) and their admixture (Admix). Pop 1 is a paraphyletic group of 115 accessions distributed mainly in countries from South Asia to the Malay Archipelago (Fig. 1c, Table S2). Pop 2 is identified as a monophyletic group with 75 accessions geographically confined to mainland Southeast Asia, mostly in Cambodia, Laos, and Vietnam, with one accession each from Myanmar and Thailand. Pop 3 is a monophyletic group of 172 accessions, mainly composed of 98 European and 34 American accessions, distributed over a wide area west of India. Two accessions each from Australia and Kenya also belong to this population. Pop 4 is a large clade of 356 accessions, with most of the East Asian accessions and some Russian or former Soviet Union accessions. Five accessions from South and East Asia, positioned as sisters to a clade comprising Pop 3 and Pop 4, showed a high degree of admixture between Pop 1 and Pop 4. We therefore considered these five accessions to be an “Admix” and excluded them from subsequent analyses, except for the core collection selection.

To quantify the degree of genetic diversity and differentiation among the four identified genetic populations, nucleotide diversity (*π*) and pairwise population fixation index (*F*_ST_) were calculated based on the raw SNPs dataset (Fig. 2). The *π* value for each population was highest in Pop 1 (1.21 ξ 10^3^), followed by Pop 3 (1.01 ξ 10^3^), Pop 2 (0.89 ξ 10^3^), and Pop 4 (0.81 ξ 10^3^). The pairwise *F*_ST_ values for each pair of populations ranged from 0.10 to 0.34, with the highest value observed between Pop 2 and Pop 4. Projecting the genetic differentiation between populations into two-dimensional space revealed a clear distinction between Pop 4 and the other three populations. Furthermore, Pop 2 and Pop 4 were differentiated from Pop 1 toward different directions in the two-dimensional space.

**Fig. 2.**
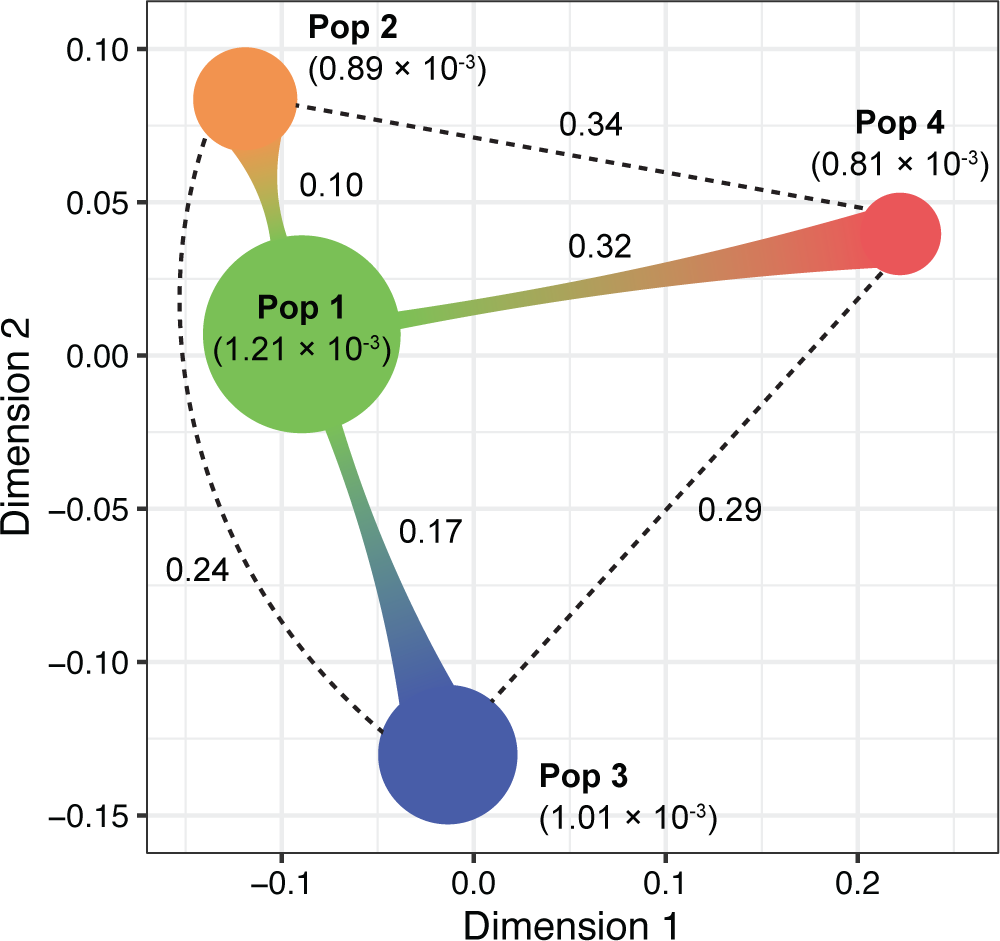
Genetic diversity and differentiation among the four genetic populations. Colored circles represent each population. The circle size reflects the nucleotide diversity (*π*) within each population, and the distance between circles represents the population fixation index (*F*_ST_) between populations. Actual values of *π* and *F*_ST_ are given in parentheses and between pairs of populations, respectively

### Associations between genetic populations and phenotypic traits

The association between the four genetic populations and phenotypic traits was investigated using the data obtained from NARO’s Plant Genetic Resources Database. For each trait, data were available for 66 to 214 of the 723 accessions used in this study (Table S3).

The quantitative traits of fruit length, fruit diameter, and seed length, differed among the populations. Pop 4 had a longer fruit length than the others (Fig. 3a), while Pop 2 had the thickest fruit diameter (Fig. 3b). The longest seed length was observed in Pop 1, while the other three populations had similar, shorter seed lengths (Fig. 3c).

**Fig. 3.**
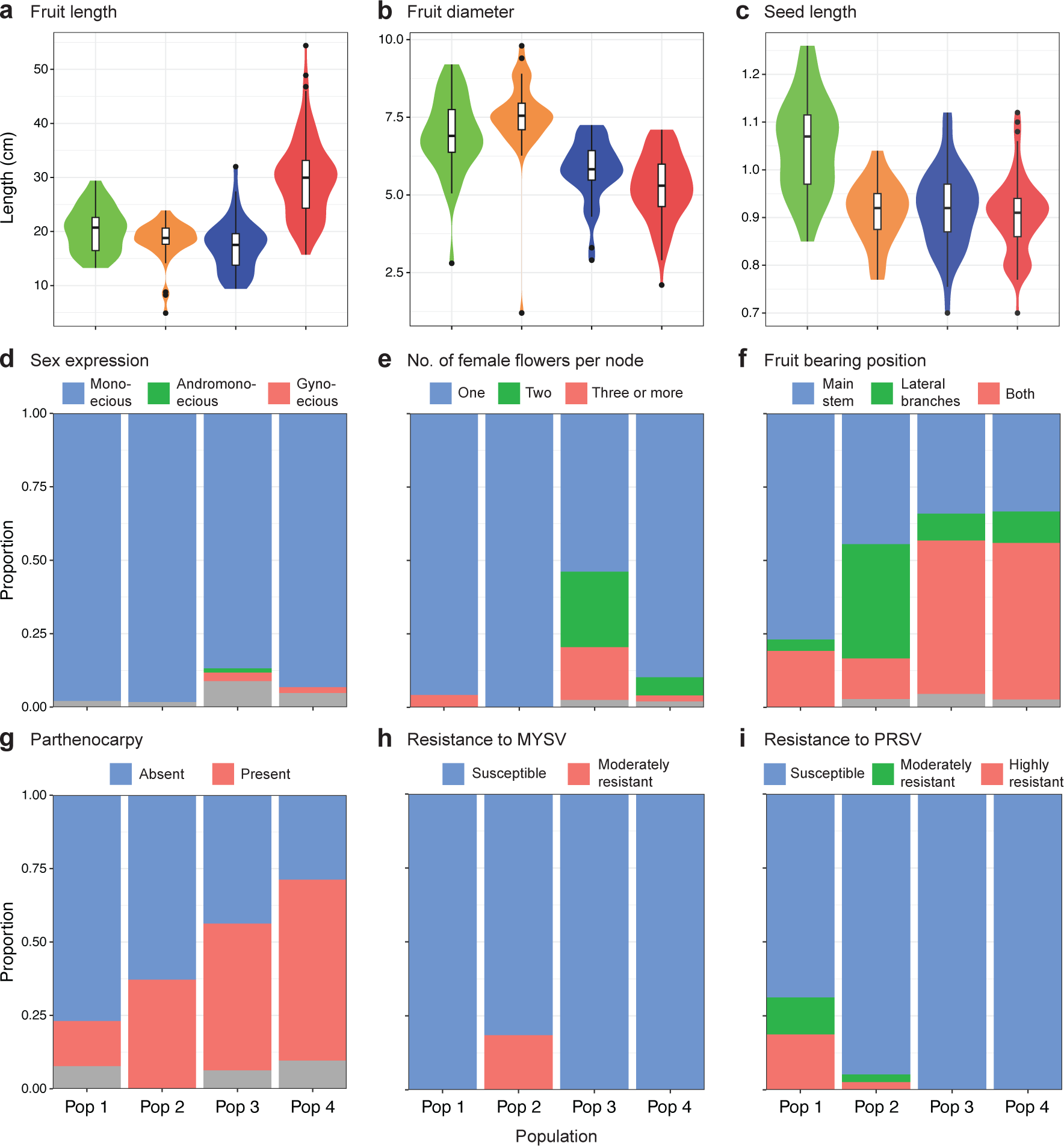
Phenotypic variation in each genetic population. **a** Fruit length (cm). **b** Fruit diameter (cm). **c** Seed length (cm). **d** Sex expression. **e** Number of female flowers per node. **f** Fruit bearing position. **g** Parthenocarpy. **h** Resistance to melon yellow spot virus (MYSV). **i** Resistance to papaya ringspot virus (PRSV). The gray color in panels **d**–**i** represents accessions that showed intra-accession segregation

Associations with the genetic populations were also inspected for yield-related traits, including sex expression, number of female flowers per node, fruit bearing position, and parthenocarpy. For sex expression, although most accessions were monoecious in all populations, Pop 3 and Pop 4 contained two and three gynoecious accessions, respectively (Fig. 3d). Pop 3 also contained one andromonoecious accession. Pop 3 had the highest proportion of accessions bearing more than one female flower per node, followed by Pop 4 (Fig. 3e). As for fruit bearing position, most of the accessions in Pop 1 fruited only on the main stem, whereas accessions fruiting on both the main stem and lateral branches were predominant in Pop 3 and Pop 4 (Fig. 3f). Pop 2 had a higher proportion of accessions fruiting only on lateral branches than the other populations. The proportion of parthenocarpic accessions increased progressively from Pop 1 to Pop 4, with more than half of the accessions showing parthenocarpy in Pop 3 and Pop 4 (Fig. 3g).

For resistance to the two viral diseases, resistant accessions were found only in Pop 1 and Pop 2. Five accessions with moderate resistance to MYSV were found exclusively in Pop 2 (Fig. 3h), and seven accessions with moderate or high resistance to PRSV were found in Pop 1 and Pop 2 (Fig. 3i).

### Subpopulation structure within the East Asian population

Traditionally, East Asian cucumbers have been classified into two types, the North China type and the South China type, each characterized by different fruit characteristics (Cui and Zhang 1991; Gebretsadik et al. 2021; Yu et al. 2022). The North China type typically produces fruits with white spines and dark green skin when immature, which turns yellow without netting at maturity. The South China type bears fruits with black spines and pale green skin when immature, turning brown with netting as it ripens. The two types also differ in their photoperiod sensitivity (Tian et al. 2021). The North China type is insensitive to photoperiod, consistently producing female flowers regardless of day length. The South China type is photoperiod-sensitive, bearing female flowers on almost every node under short-day conditions, but with significant suppression of female flower development under long-day conditions.

Our phylogenetic analysis identified two large subclades (Subclades 1 and 2) within Pop 4, comprising 144 and 167 accessions, respectively (Fig. 1a). To examine the correspondence between this subpopulation structure and the traditional classification of East Asian cucumbers, we compared mature fruit skin color, mature fruit skin netting, and fruit spine color between accessions belonging to the two subclades. The results showed differences between the subclades in all the traits examined (Fig. 4a). For mature fruit skin color, 14 (54%) of 26 accessions in Subclade 1 for which trait data were available on NARO’s database had yellow skin, whereas 28 (48%) of 58 accessions in Subclade 2 had brown skin. Similarly, for the presence of netting on mature fruit skin, only 3 (9%) of 34 accessions in Subclade 1 exhibited netting, compared to 30 (63%) of 48 accessions in Subclade 2. Fruit spine color followed a similar pattern: 19 (73%) of 26 accessions in Subclade 1 had white spines, while 32 (54%) of 59 accessions in Subclade 2 had black spines. Taken together, these results support the correspondence between the two subclades within Pop 4 and the traditional classification of East Asian cucumbers into the North and South China types.

**Fig. 4.**
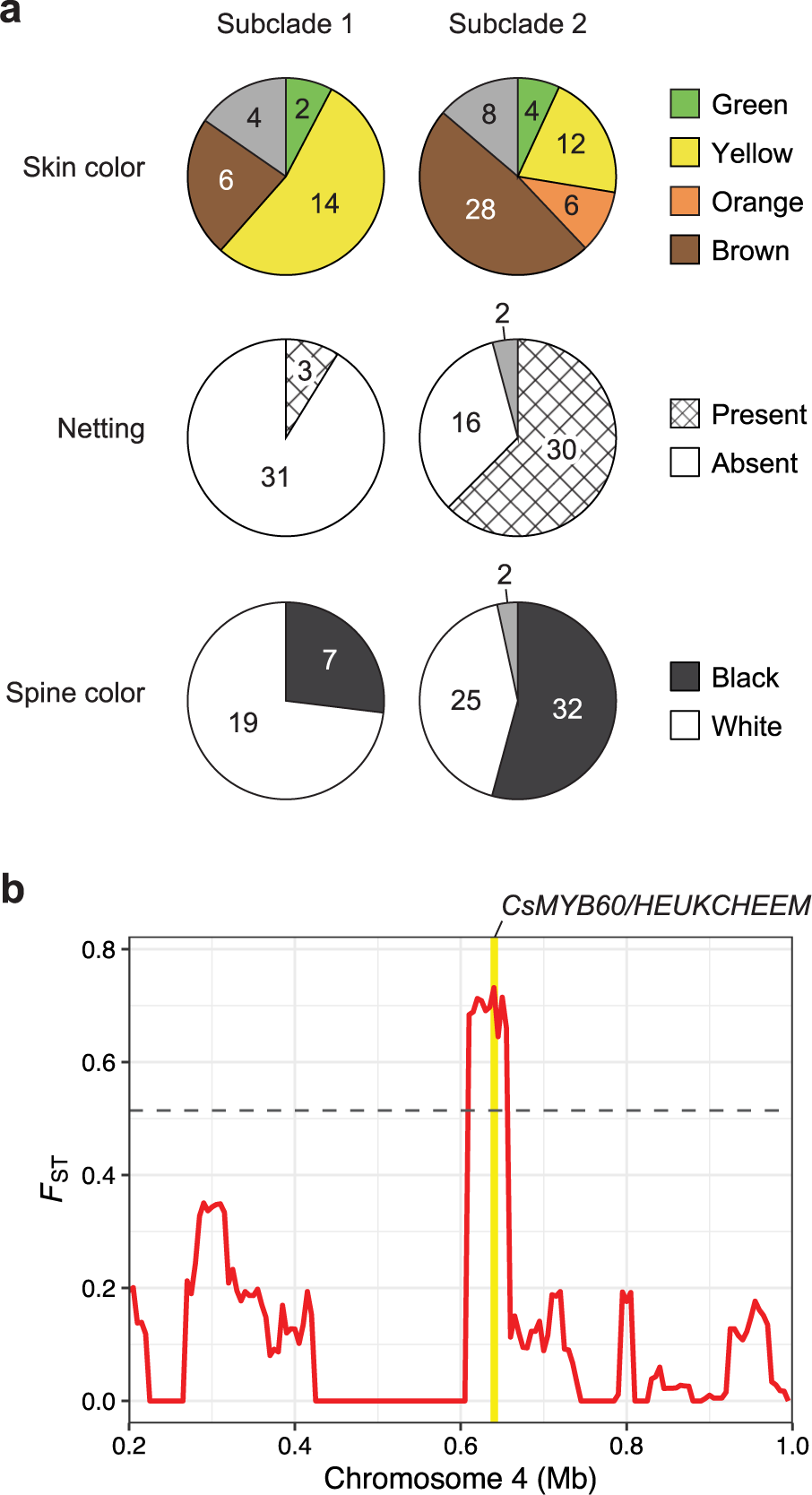
Differences in fruit traits and corresponding local genetic differentiation between two subclades within the East Asian population. **a** Number of accessions with different mature fruit skin color, netting, and spine color in each subclade. The gray color in each pie represents accessions that showed intra-accession segregation. **b** Fixation index (*F*_ST_) between the two subclades surrounding the *CsMYB60/HEUKCHEEM* gene responsible for fruit spine color. The position of the gene is highlighted in yellow. The horizontal dashed line indicates the genome-wide threshold for the top 1% highly divergent region

A series of genetic studies has identified that fruit spine color in cucumber is controlled by *CsMYB60/HEUKCHEEM* (*CsaV3_4G001130*) through the transcriptional regulation of anthocyanin biosynthesis-related genes (Li et al. 2013; Liu et al. 2019b; Zhang et al. 2019). To investigate the association of *CsMYB60/HEUKCHEEM* with the observed difference in fruit spine color, we calculated local *F*_ST_ values between the two subclades for 50-kb sliding windows with a step size of 5 kb. The result showed that the genomic region surrounding *CsMYB60/HEUKCHEEM* had exceptionally high *F*_ST_ values above the top 1% genome-wide threshold (0.514), indicating significant differentiation between the two subclades in this genomic region (Fig. 4b).

### Development of the World Cucumber Core Collection

To objectively select candidates for the core collection, 621 accessions, excluding the outgroup and F_1_ accessions, were ranked to maximize the cumulative allelic coverage of the filtered SNPs, using GenoCore (Jeong et al. 2017). The result showed that the 16, 28, 51, 111, and 512 top-ranked accessions represent 80%, 85%, 90%, 95%, and 100% of alleles of the filtered 61,367 SNPs, respectively (Fig. 5a). Balancing the allelic coverage and collection size, we selected a total of 100 accessions for the core collection, designated as the World Cucumber Core Collection (WCC). The WCC consists of 79 accessions representing 93.2% of the alleles, and 21 accessions manually selected for their historical significance, geographic origin, or unique traits such as disease resistance, hermaphrodite, and orange flesh color. The accessions selected for the WCC are listed in Table S4.

**Fig. 5.**
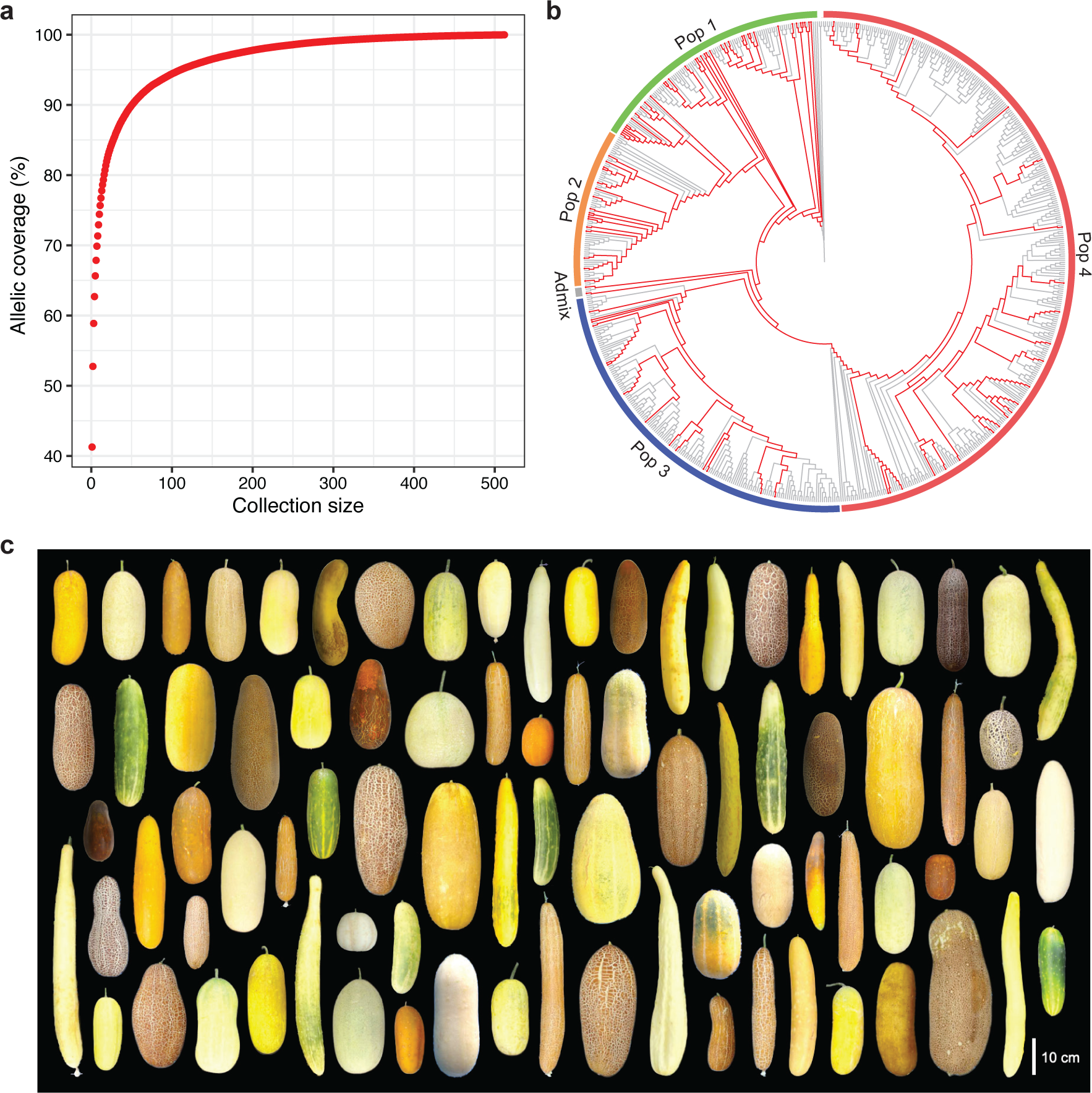
Development of the World Cucumber Core Collection (WCC). **a** Allelic coverage of the 61,367 filtered SNPs versus collection size selected by GenoCore. **b** The same phylogeny as in Fig. 1a, showing only topology. Accessions selected for the WCC are highlighted in red. **c** Fruit morphological diversity of the WCC (partial)

The representativeness of the WCC was verified by several observations. The accessions selected were widely distributed throughout the phylogeny (Fig. 5b), covering all identified genetic populations and broad geographic regions (Table S2). The mature fruit morphology of the WCC showed remarkable diversity in various traits, such as size, shape, skin color, and pattern (Fig. 5c). These results confirmed that the WCC represents a significant portion of the genetic diversity present in the entire collection used in this study.

## Discussion

While classical studies based on dozens of isozyme loci or RAPD markers had claimed a narrow genetic diversity of cucumber (Horejsi and Staub 1999; Meglic et al. 1996; Meglic and Staub 1996; Staub et al. 1999; Staub et al. 1997), more recent studies based on thousands or millions of SNPs have revealed clear genetic differentiation between geographic regions (Lv et al. 2012; Qi et al. 2013; Wang et al. 2018). Our analyses based on the genome-wide SNPs also identified four genetic populations (Pops 1–4) highly associated with the geographic distribution (Fig. 1). Pop 1 was mainly distributed in South Asia and was positioned deepest in the phylogenetic tree. The inclusion of the wild cucumber (*C. s*. var. *hardwickii*) accession and the highest nucleotide diversity (Fig. 2) in this population supported the South Asian origin of cucumber, and also suggested an identity between this population and the Indian group reported in previous studies (Lv et al. 2012; Qi et al. 2013; Wang et al. 2018). The geographic distribution of Pop 2 was highly restricted in mainland Southeast Asia, especially in Cambodia, Laos, and Vietnam, which are in close proximity to Xishuangbanna, where an endemic population has been reported in previous studies (Liu et al. 2019a; Qi et al. 2013). The taxonomic description of *C*. *s*. var. *xishuangbannanensis* states that this variety is distributed not only in Xishuangbanna but also in Laos, Vietnam, and probably Myanmar (Renner 2017), suggesting a possible identity between Pop 2 and the Xishuangbanna group in previous studies. Phenotypically, Pop 2 tended to have thicker fruits than the other populations (Fig. 3b) and included accessions with orange flesh (Shimomura et al. 2016), which is in line with the characteristics of *C*. *s*. var. *xishuangbannanensis*. However, since our material did not include a single accession from Xishuangbanna, further studies using both Pop 2 and Xishuangbanna accessions would be needed to verify the identity of these two populations. Pop 3 and Pop 4 are thought to be the two cultivated populations that dispersed westward and eastward from South Asia, respectively. Pop 3 is distributed over a wide area west of India, including Europe, the Americas, and West and Central Asia, and would correspond to the Eurasian group in previous studies. Pop 4 has a restricted distribution in East Asia, with some occurrence in Russia or the former Soviet Union, which would correspond to the East Asian group in previous studies. The five Admix accessions were likely of hybrid origin between Pop 1 and Pop 4, based on their intermediate position in the phylogenetic tree and the composition of the genetic admixture (Fig. 1a, b). Overall, our results largely supported the genetic–geographic population structure of cucumber reported in previous studies (Liu et al. 2019a; Lv et al. 2012; Qi et al. 2013; Wang et al. 2018), with a potential update on the distribution range of the Xishuangbanna group.

The genetic diversity of each population, as quantified by *π*, was highest in Pop 1 and lower in the other three populations (Fig. 2), supporting the genetic bottleneck that natural/artificial selection in each geographic region resulted in reduced genetic diversity relative to the ancestral population (Qi et al. 2013). Genetic differentiation among the populations quantified by pairwise *F*_ST_ shows that Pop 4 stands apart from the other three populations (Fig. 2), which is consistent with the result of the population structure analysis at *K* = 2 that distinguished Pop 4 from all other accessions (Fig. 1b). The distinctiveness of the East Asian population was also reported by Wang et al. (2018), suggesting that independent and intensive selection has been carried out in East Asia. Notably, despite their geographical proximity, the highest genetic differentiation was observed between Pop 2 and Pop 4, and these two populations differentiated toward different directions from Pop 1 in two-dimensional space (Fig. 2). These results imply that Pop 2 and Pop 4 arrived at their respective regions via independent routes during the eastward dispersal from South Asia, rather than following a common route along the Southeast Asian coastline. Similar hypotheses have been proposed for melon (*C. melo*), another important crop in the genus *Cucumis*, given the clear genetic differences between landraces in Southeast and East Asia (Duong et al. 2021; Naznin et al. 2024; Nhi et al. 2010). This hints at the possibility that the two cucurbit crops partially share their dispersal routes, especially to Southeast and East Asia.

Although phenotypic trait data were available for only a limited number of accessions on the database, different patterns were observed among the populations. Fruit shape was longer in Pop 4 (Fig. 3a) and thicker in Pop 2 (Fig. 3b), suggesting that different fruit morphologies may have been preferred and selected in each region. Seed size was larger in Pop 1 than in the other populations (Fig. 3c). This may reflect a preference for less seedy fruits that do not interfere with the crunchy texture of cucumbers, which are mostly eaten without removing the seeds. Sex expression, number of female flowers per node, fruit bearing position, and parthenocarpy are considered to be yield-related traits because they are directly related to how many fruits can be set on a plant. Parthenocarpy may also be desirable as it produces seedless fruits. For all these traits, Pop 3 and Pop 4 tended to contain more accessions with traits favoring high yield (Fig. 3d–g). In contrast, for resistance to the two viral diseases, only Pop 1 and Pop 2 contained resistant accessions (Fig. 3h, i), reaffirming the crucial value of these populations as breeding material for disease resistance.

Traditionally, East Asian cucumbers have been classified into the North and South China types (Cui and Zhang 1991; Gebretsadik et al. 2021; Yu et al. 2022). It is believed that the two types were introduced to China through independent routes and subsequently spread to other countries in East Asia. The history of the introduction of the North China type is well documented in the Chinese encyclopedic herbal; it was introduced from the west via the Silk Road by the diplomat Zhang Qian (164–114 BC) during the Han Dynasty (Li 1590). On the other hand, no written records have been found on the introduction of the South China type, but it has been hypothesized that this type of cucumber was introduced directly from the south through ancient Himalayan trade routes around the border with Myanmar.

Of the 723 cucumber accessions used in this study, 352 were from East Asia, mainly from China and Japan (Fig. 1c, Table S2). This number of East Asian accessions allowed us to further investigate the subpopulation structure within Pop 4, the East Asian population. Our analysis revealed that the three fruit traits which typically distinguish the North and South China types also differ between the two subclades within Pop 4 (Fig. 4a). Specifically, Subclade 1 tended to have fruits with yellow skins, no netting, and white spines, while Subclade 2 had fruits with brown skins, netting, and black spines, consistent with the characteristics of the North and South China types, respectively. Furthermore, corresponding to the difference in fruit spine color, the genomic region surrounding *CsMYB60/HEUKCHEEM*, the gene cloned to be responsible for fruit spine color (Liu et al. 2019b; Zhang et al. 2019), showed significant differentiation between the two subclades (Fig. 4b). These results provided genetic support for the traditional classification of East Asian cucumbers and their correspondence to the two subclades identified within the East Asian population.

With a holistic approach to take into account genetic diversity, historical significance, geographic origin, and unique traits, we developed our WCC consisting of 100 accessions. This core collection represents at least 93.2% of the genetic diversity present in the 723 accessions used in this study, and includes several accessions with resistance to major fungal, bacterial, and viral diseases (Table S4). The representativeness of the WCC was supported by the inclusion of accessions from all identified genetic populations and diverse geographic origins (Table S2), their widespread distribution throughout the phylogeny (Fig. 5b), and the remarkable diversity in fruit morphology (Fig. 5c). Compared to the core collections developed in previous studies (Lv et al. 2012; Staub et al. 2002; Wang et al. 2018), the WCC has a relatively small collection size while featuring a significant portion of the genetic diversity. This compactness, coupled with its ability to compensate for the underrepresentation of Southeast Asian accessions in the prior core collections, will provide a unique advantage to fit into a broader range of studies.

The WCC will be publicly available from the NARO Genebank (https://www.gene.affrc.go.jp/databases-core_collections_en.php) to researchers and breeders worldwide, upon request. The phylogenetic relationships and population structure, their association with geographic distribution and phenotypic traits, and the core collection presented in this study will be valuable resources to elucidate the dispersal history of cucumber and facilitate the efficient use and management of genetic resources in cucumber research and breeding.

## Statements and Declarations

### Acknowledgments

We thank Prof. Dr. Koichiro Ushijima (Okayama University, Japan) for his helpful advice on GBS library preparation.

### Author contribution statement

KK, NT, and YK conceived the project. MS and KS provided materials. GS, TPD, MNP, TTD, and ONI performed the experiments. GS analyzed the data, prepared figures, and drafted the manuscript. YM, HN, KT, and KK provided advice on the experimental implementation and reviewed the manuscript.

### Data availability

The raw GBS reads for each accession are available at the DDBJ Sequence Read Archive (DRA) under the BioProject PRJDB14335.

## References

Alexander DH, Novembre J, Lange K (2009) Fast model-based estimation of ancestry in unrelated individuals. Genome Res 19:1655–1664. 10.1101%2Fgr.094052.109

Behr AA, Liu KZ, Liu-Fang G, et al (2016) pong: fast analysis and visualization of latent clusters in population genetic data. Bioinformatics 32:2817–2823. 10.1093/bioinformatics/btw327

Bisht IS, Bhat KV, Tanwar SPS, et al (2004) Distribution and genetic diversity of *Cucumis sativus* var. *hardwickii* (Royle) Alef in India. J Hortic Sci Biotechnol 79:783–791. 10.1080/14620316.2004.11511843

Chen S, Zhou Y, Chen Y, Gu J (2018) fastp: an ultra-fast all-in-one FASTQ preprocessor. Bioinformatics 34:i884–i890. 10.1093/bioinformatics/bty560

Chomicki G, Schaefer H, Renner SS (2020) Origin and domestication of Cucurbitaceae crops: insights from phylogenies, genomics and archaeology. New Phytol 226:1240–1255. 10.1111/nph.16015

Cui H, Zhang X (1991) Cucumber cultivar improvement in the People’s Republic of China. Rep Cucurbit Genet Coop 14:5–7.

Danecek P, Auton A, Abecasis G, et al (2011) The variant call format and VCFtools. Bioinformatics 27:2156–2158. 10.1093%2Fbioinformatics%2Fbtr330

Danecek P, Bonfield JK, Liddle J, et al (2021) Twelve years of SAMtools and BCFtools. Gigascience 10:giab008. 10.1093/gigascience/giab008

Duong TT, Dung TP, Tanaka K, et al (2021) Distribution of two groups of melon landraces and inter-group hybridization enhanced genetic diversity in Vietnam. Breed Sci 71:564–574. 10.1270/jsbbs.20090

Elshire RJ, Glaubitz JC, Sun Q, et al (2011) A robust, simple genotyping-by-sequencing (GBS) approach for high diversity species. PLoS One 6:e19379. 10.1371/journal.pone.0019379

Endl J, Achigan-Dako EG, Pandey AK, et al (2018) Repeated domestication of melon (*Cucumis melo*) in Africa and Asia and a new close relative from India. Am J Bot 105:1662–1671. 10.1002/ajb2.1172

Frankel OH, Brown AHD (1984) Plant genetic resources today: a critical appraisal. In: Holden JHW, Williams JT (eds) Crop genetic resources: conservation and evaluation. George Allan and Unwin, London, pp 249–257

Gebretsadik K, Qiu X, Dong S, et al (2021) Molecular research progress and improvement approach of fruit quality traits in cucumber. Theor Appl Genet 134:3535–3552. 10.1007/s00122-021-03895-y

Hoang DT, Chernomor O, von Haeseler A, et al (2018) UFBoot2: improving the ultrafast bootstrap approximation. Mol Biol Evol 35:518–522. 10.1093/molbev/msx281

Horejsi T, Staub JE (1999) Genetic variation in cucumber (*Cucumis sativus* L.) as assessed by random amplified polymorphic DNA. Genet Resour Crop Evol 46:337–350. 10.1023/A:1008650509966

Jeong S, Kim J-Y, Jeong S-C, et al (2017) GenoCore: A simple and fast algorithm for core subset selection from large genotype datasets. PLoS One 12:e0181420. 10.1371/journal.pone.0181420

Kawazu Y, Kato M, Tran TTH, Nguyen VK (2017) Collaborative exploration of plant genetic resources in Vietnam, 2016. Annual Report on Exploration and Introduction of Plant Genetic Resources 33:89–114. 10.24514/00001097

Kawazu Y, Kuzuya M, Ouch S, et al (2020) Collaborative exploration of Cucurbitaceae genetic resources in eastern Cambodia, 2019. Annual Report on Exploration and Introduction of Plant Genetic Resources 36:92–111. 10.24514/00005677

Knerr LD, Staub JE (1992) Inheritance and linkage relationships of isozyme loci in cucumber (*Cucumis sativus* L.). Theor Appl Genet 84:217–224. 10.1007/bf00224003

Knerr LD, Staub JE, Holder DJ, May BP (1989) Genetic diversity in *Cucumis sativus* L. assessed by variation at 18 allozyme coding loci. Theor Appl Genet 78:119–128. 10.1007/bf00299764

Lee H-Y, Kim J-G, Kang B-C, Song K (2020) Assessment of the genetic diversity of the breeding lines and a genome wide association study of three horticultural traits using worldwide cucumber (*Cucumis* spp.) germplasm collection. Agronomy 10:1736. 10.3390/agronomy10111736

Lewis PO (2001) A likelihood approach to estimating phylogeny from discrete morphological character data. Syst Biol 50:913–925. 10.1080/106351501753462876

Li H (2013) Aligning sequence reads, clone sequences and assembly contigs with BWA-MEM. arXiv 1303.3997. 10.48550/arXiv.1303.3997

Li H, Wang S, Chai S, et al (2022) Graph-based pan-genome reveals structural and sequence variations related to agronomic traits and domestication in cucumber. Nat Commun 13:682. 10.1038/s41467-022-28362-0

Li H-L (1969) The vegetables of ancient China. Econ Bot 23:253–260.

Li Q, Li H, Huang W, et al (2019) A chromosome-scale genome assembly of cucumber (*Cucumis sativus* L.). Gigascience 8. 10.1093/gigascience/giz072

Li S (1590) Vegetables, part III: cucurbits. In: Compendium of Materia Medica, China, pp 14–15

Li Y, Wen C, Weng Y (2013) Fine mapping of the pleiotropic locus *B* for black spine and orange mature fruit color in cucumber identifies a 50 kb region containing a R2R3-MYB transcription factor. Theor Appl Genet 126:2187–2196. 10.1007/s00122-013-2128-3

Liu B, Guan D, Zhai X, et al (2019a) Selection footprints reflect genomic changes associated with breeding efforts in 56 cucumber inbred lines. Hortic Res 6:127. 10.1038/s41438-019-0209-4

Liu M, Zhang C, Duan L, et al (2019b) *CsMYB60* is a key regulator of flavonols and proanthocyanidans that determine the colour of fruit spines in cucumber. J Exp Bot 70:69–84. 10.1093/jxb/ery336

Lv J, Qi J, Shi Q, et al (2012) Genetic diversity and population structure of cucumber (*Cucumis sativus* L.). PLoS One 7:e46919. 10.1371/journal.pone.0046919

Matsunaga H, Matsushima K, Tanaka K, et al (2015) Collaborative exploration of the Solanaceae and Cucurbitaceae vegetable genetic resources in Cambodia, 2014. Annual Report on Exploration and Introduction of Plant Genetic Resources 31:169–187. 10.24514/00004732

Meglic V, Serquen F, Staub JE (1996) Genetic diversity in cucumber (*Cucumis sativus* L.): I. A reevaluation of the U.S. germplasm collection. Genet Resour Crop Evol 43:533–546. 10.1007/BF00138830

Meglic V, Staub JE (1996) Genetic diversity in cucumber (*Cucumis sativus* L.): II. An evaluation of selected cultivars released between 1846 and 1978. Genet Resour Crop Evol 43:547–558. 10.1007/BF00138831

Minh BQ, Schmidt HA, Chernomor O, et al (2020) IQ-TREE 2: new models and efficient methods for phylogenetic inference in the genomic era. Mol Biol Evol 37:1530– 1534. 10.1093/molbev/msaa015

Murray MG, Thompson WF (1980) Rapid isolation of high molecular weight plant DNA. Nucleic Acids Res 8:4321–4325. 10.1093/nar/8.19.4321

Naznin PM, Imoh ON, Tanaka K, et al (2024) Analysis of genetic diversity and population structure in Cambodian melon landraces using molecular markers. Genet Resour Crop Evol 71:1067–1083. 10.1007/s10722-023-01677-7

Nei M, Li WH (1979) Mathematical model for studying genetic variation in terms of restriction endonucleases. Proc Natl Acad Sci U S A 76:5269–5273. 10.1073/pnas.76.10.5269

Nhi PTP, Akashi Y, Hang TTM, et al (2010) Genetic diversity in Vietnamese melon landraces revealed by the analyses of morphological traits and nuclear and cytoplasmic molecular markers. Breed Sci 60:255–266. 10.1270/jsbbs.60.255

Okuizumi H, Phengphachanh B, Hongphakdy K, et al (2015) Collaborative exploration for plant genetic resources in Laos, December, 2014. Annual Report on Exploration and Introduction of Plant Genetic Resources 31:225–293. 10.24514/00004735

Paris HS, Daunay MC, Janick J (2012) Occidental diffusion of cucumber (*Cucumis sativus*) 500-1300 CE: two routes to Europe. Ann Bot 109:117–126. 10.1093/aob/mcr281

Qi C, Yuan Z, Li Y (1983) A new type of cucumber - *Cucumis sativus* L. var. *xishuangbannanesis*. Acta Horticulturae Sinica 10:259–263.

Qi J, Liu X, Shen D, et al (2013) A genomic variation map provides insights into the genetic basis of cucumber domestication and diversity. Nat Genet 45:1510–1515. 10.1038/ng.2801

Renner SS (2017) A valid name for the Xishuangbanna gourd, a cucumber with carotene-rich fruits. PhytoKeys:87–94. 10.3897/phytokeys.85.17371

Renner SS, Schaefer H, Kocyan A (2007) Phylogenetics of *Cucumis* (Cucurbitaceae): Cucumber (*C. sativus*) belongs in an Asian/Australian clade far from melon (*C. melo*). BMC Evol Biol 7:58. 10.1186/1471-2148-7-58

Sebastian P, Schaefer H, Telford IRH, Renner SS (2010) Cucumber (*Cucumis sativus*) and melon (*C. melo*) have numerous wild relatives in Asia and Australia, and the sister species of melon is from Australia. Proc Natl Acad Sci U S A 107:14269–14273. 10.1073/pnas.1005338107

Shigita G, Dung TP, Pervin MN, et al (2023) Elucidation of genetic variation and population structure of melon genetic resources in the NARO Genebank, and construction of the World Melon Core Collection. Breed Sci 73:269–277. 10.1270/jsbbs.22071

Shimomura K, Ohm MS, San Thein M (2020) Collaborative exploration of Cucurbitaceae vegetable genetic resources in Myanmar in 2019. Annual Report on Exploration and Introduction of Plant Genetic Resources 36:148–158. 10.24514/00005680

Shimomura K, Sugiyama K, Yoshioka Y, et al (2016) Collaborative exploration of plant genetic resources in Vietnam, 2015. Annual Report on Exploration and Introduction of Plant Genetic Resources 32. 10.24514/00004665

Staub JE, Dane F, Reitsma K, et al (2002) The formation of test arrays and a core collection in cucumber using phenotypic and molecular marker data. J Am Soc Hortic Sci 127:558–567. 10.21273/JASHS.127.4.558

Staub JE, Serquen FC, Horejsi T, Chen J-f (1999) Genetic diversity in cucumber (*Cucumis sativus* L.): IV. An evaluation of Chinese germplasm. Genet Resour Crop Evol 46:297–310. 10.1023/A:1008663225896

Staub JE, Serquen FC, McCreight JD (1997) Genetic diversity in cucumber (*Cucumis sativus* L.): III. An evaluation of Indian germplasm. Genet Resour Crop Evol 44:315–326. 10.1023/A:1008639103328

Sugiyama M, Ebana K, Kami D, et al (2015) Collaborative exploration of Cucurbitaceous crops in Vietnam, 2014. Annual Report on Exploration and Introduction of Plant Genetic Resources 31:189–201. 10.24514/00004733

Tanaka K, Duong TT, Yamashita H, et al (2016) Collection of cucurbit crops (Cucurbitaceae) from eastern Cambodia, 2015. Annual Report on Exploration and Introduction of Plant Genetic Resources 32:109–137. 10.24514/00004658

Tanaka K, Shigita G, Dung TP, et al (2019) Collection of melon and other cucurbitaceous crops in Cambodia in 2017. Annual Report on Exploration and Introduction of Plant Genetic Resources 35:121–146. 10.24514/00003226

Tanaka K, Shigita G, Sophea Y, et al (2017) Collection of melon and other cucurbitaceous crops in Cambodia in 2016. Annual Report on Exploration and Introduction of Plant Genetic Resources 33:175–205. 10.24514/00001100

Tian Z, Jahn M, Qin X, et al (2021) Genetic and transcriptomic analysis reveal the molecular basis of photoperiod-regulated flowering in Xishuangbanna cucumber (*Cucumis sativus* L. var. *xishuangbannesis* Qi et Yuan). Genes (Basel) 12:1064. 10.3390/genes12071064

Wang X, Bao K, Reddy UK, et al (2018) The USDA cucumber (*Cucumis sativus* L.) collection: genetic diversity, population structure, genome-wide association studies, and core collection development. Hortic Res 5:64. 10.1038/s41438-018-0080-8

Weir BS, Cockerham CC (1984) Estimating *F*-statistics for the analysis of population structure. Evolution 38:1358–1370. 10.1111/j.1558-5646.1984.tb05657.x

Wickland DP, Battu G, Hudson KA, et al (2017) A comparison of genotyping-by-sequencing analysis methods on low-coverage crop datasets shows advantages of a new workflow, GB-eaSy. BMC Bioinformatics 18:586. 10.1186/s12859-017-2000-6

Yashiro K, Tanaka K, Yon S, et al (2019) Collaborative exploration of Cucurbitaceae vegetable genetic resources in western and northwestern Cambodia in 2018. Annual Report on Exploration and Introduction of Plant Genetic Resources 35:147–161. 10.24514/00003227

Yu B, Ming F, Liang Y, et al (2022) Heat stress resistance mechanisms of two cucumber varieties from different regions. Int J Mol Sci 23:1817. 10.3390%2Fijms23031817

Zhang C, Win KT, Kim Y-C, Lee S (2019) Two types of mutations in the *HEUKCHEEM* gene functioning in cucumber spine color development can be used as signatures for cucumber domestication. Planta 250:1491–1504. 10.1007/s00425-019-03244-w

