## Supplemental Figures for "Genetic characterization of cucumber genetic resources in the NARO Genebank indicates their multiple dispersal trajectories to the East"

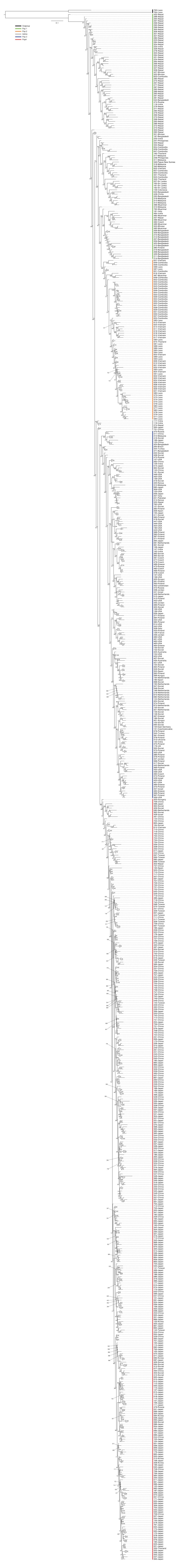


**Fig. S1** Maximum-likelihood phylogenetic tree of the 723 cucumber accessions. Tip labels indicate the sample ID and country of origin shown in Supplemental Table 1, separated by a dash. Vertical line next to the tip labels color-codes the genetic population to which each sample belongs. Support values assessed from 1,000 ultra bootstrap replicates are given at the nodes

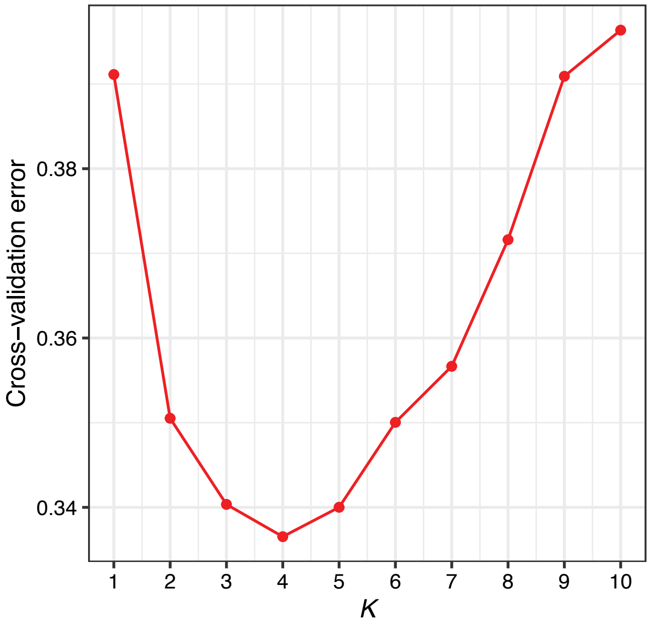


**Fig. S2** Cross-validation errors for different values of assumed ancestral populations (*K*) from 1 to 10 in the ADMIXTURE analysis
